# Logic Gate Activated Lysosome Targeting DNA Nanodevice for Controlled Proteins Degradation

**DOI:** 10.1101/2023.08.29.555427

**Authors:** Yuzhe Shang, Longyi Zhu, Yang Xiao, Songyuan Du, Ruoyang Ji, Bin Li, Jialiang Chen, Shengyuan Deng, Kewei Ren

**Author notes:** These authors contributed equally to this work (Yuzhe Shang, Longyi Zhu). Supporting information for this article is given via a link at the end of the document.

## Abstract

Targeted protein degradation (TPD) is a powerful technique for regulation of protein homeostasis. Current TPD mainly focuses on the therapeutical consequences rather than the operation processes of the molecular tools. Herein, we construct a platform for precisely manipulate the protein degradation by activatable lysosome targeting DNA nanodevices. In the design, a lysosome-targeting CD63 aptamer is locked by the single-stranded DNA with a photocleavable group and a disulfide bond. This locked CD63 aptamer is connected with the aptamer targeting the protein of interest via double-stranded DNA linkages to form the logic-gate activated lysosome targeting DNA nanodevice (LALTD). With the UV light and endogenous GSH as inputs, AND logic-gate is built to efficiently manipulate the protein delivery processes by LALTD. The protein of interest could be degraded via efficient lysosome hydrolysis. Further studies shows that the logic-gate operation could be used for modulating the T cell-mediated antitumor immunity. The modularly designed activatable lysosome targeting DNA nanodevices exhibits good stability, controllability, programmability and universality, providing a new prospect for accurate protein degradation and precise therapy.

**Entry for the Table of Contents:** Through rational integration of dual molecular switches with bispecific aptamer systems, a logic-gate activated lysosome targeting DNA nanodevice (LALTD) was developed for precisely controlled process of protein hydrolysis in living cells. The designed LALTD system provide a general platform for designing accurate protein degradation.

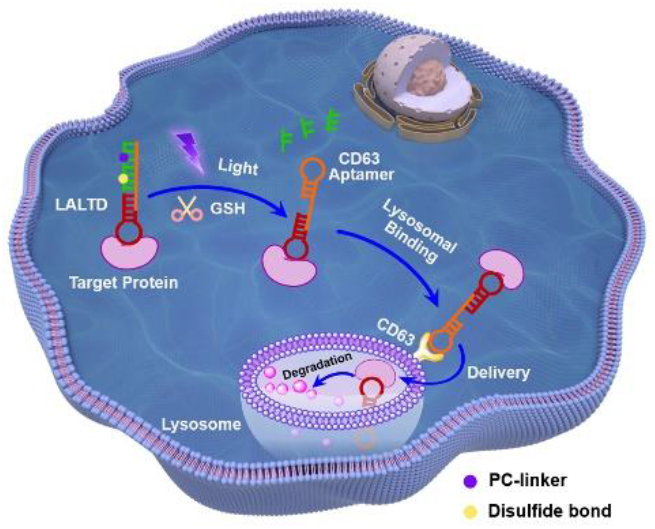

---

The removal of abnormal and pathogenic proteins is crucial in regulating cellular proteostasis.^[1]^ Traditional protein eliminating methods are based on genetic engineering and inhibitor screening.^[2,3]^ Recently, targeted protein degradation (TPD) shows broader scope of application covering “undruggable” targets and has less time-costs, which becomes a powerful tool for designing therapeutic strategies.^[4,5]^ TPD techniques hijack the protein hydrolysis pathways to eliminate the protein of interest (POI) by using different types of synthetical degraders, such as molecular glues,^[6]^ proteolysis targeting chimeras (PROTACs),^[7]^ lysosome-targeting chimeras (LYTACs)^[8]^ and autophagy-targeting chimera (AUTAC)^[9]^ etc. By customization of ligands on the degrader for both POI and the functional protein in the hydrolysis pathways, many excellent works on TPD have been continuously reported.^[10-12]^ However, current TPD techniques still face many challenges. One of the concerned issues is the risk of possible off-target effects.^[13]^ Introduction of aptamer moieties offers an attractive approach to improve the cell targeting ability of the degrader.^[14]^ But the evaluation on the efficiency of degradation process after recognition of POI still remains largely underdeveloped. Therefore, a universal system for precise manipulation of protein degradation is highly desirable.

**Scheme 1.**
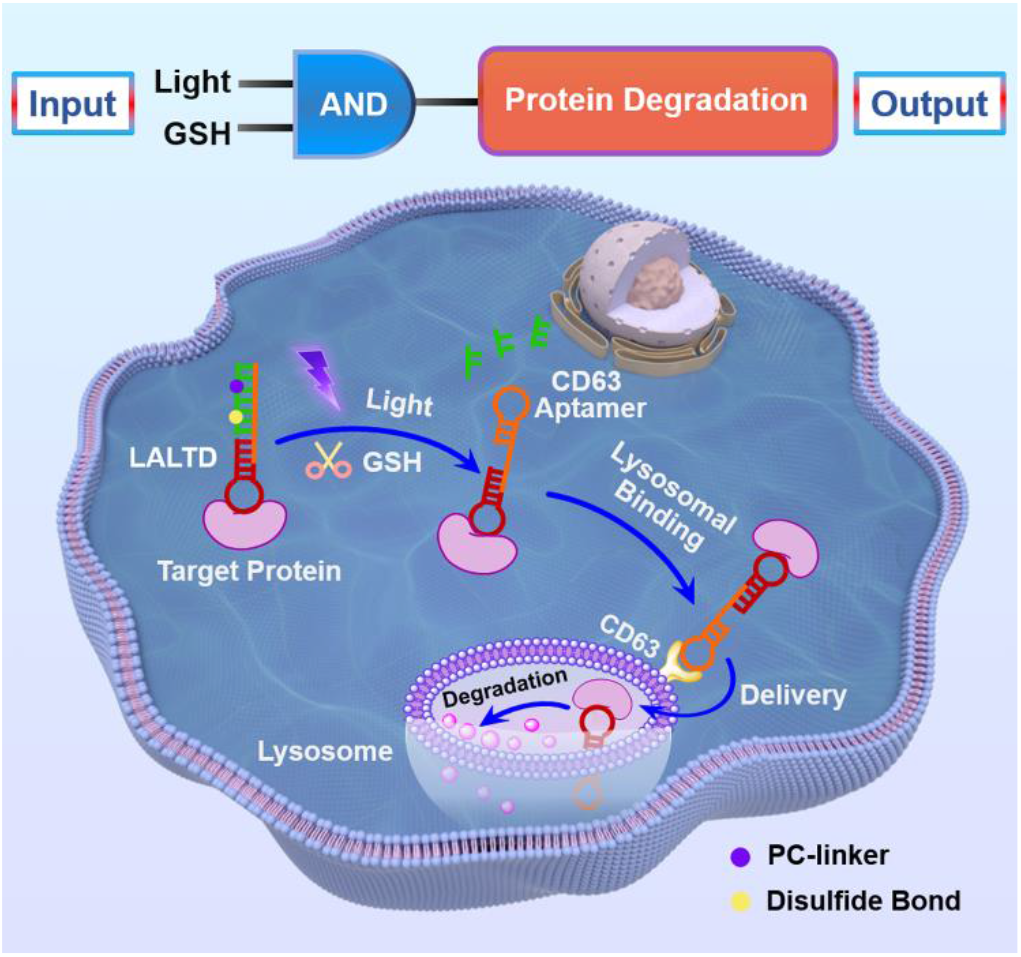
Schematic illustration of logic-gate activated lysosome targeting DNA nanodevice for controlled protein degradation.

DNA nanostructures have been widely applied to design nanodevices for theragnostic proposes.^[15,16]^ The programmable base-pairings, easy post-modification and good biocompatibility could further promote DNA nanodevices to accomplish intelligent operations in the biosystems, such as catalytic amplifiers,^[17]^ molecular walkers^[18]^ and logic gates.^[19]^ By designing operations according to the specific biomolecular cue, DNA-based logic gates are particularly useful for manipulating the bioprocesses *in vivo*.^[20-22]^ For examples, a dual-responsive DNAzyme logic system only responds to both high concentration of miR-21 and H2O2 to allow cell-specific chemodynamic therapy.^[23]^ The “AND” gate nanodevice for both protons and ATP inputs was reported for successfully modulating the cell surface receptor assembly and down-stream signaling.^[24]^ The multivariate recognitions from inputs could increase the specificity of the nanodevices and decrease the potential side effects.^[25,26]^ Thus, we envision that incorporating DNA-based logic gates to the protein degradation process will help to improve the therapeutic efficiency.

Herein, we present a series of activatable lysosome targeting DNA nanodevice for controlled protein degradation in living cells. Through integration of molecular switches with bispecific aptamer chimeras, photoactivated lysosome targeting DNA nanodevice (PALTD) is developed for UV-light triggered protein degradation *via* lysosome hydrolysis. Further introduction of the disulfide-bond domain with PALTD achieves logic-gate activated lysosome targeting DNA nanodevice (LALTD), which allows both UV light and endogenous GSH as logic manipulation for controlled protein degradation in cancer cells (Scheme 1). In addition, we have successfully verified the LALTD systems in T cell-mediated antitumor immunity and different POI recognition scenarios, demonstrating a promising general platform for precise protein degradation.

In order to design PALTD system, responsive lysosome targeting moieties were initially designed. We used CD63 as the specific receptor to mediate lysosomal targeting, internalization and protein delivery, which is a member of the four span protein superfamily and widely expressed on the lysosomes. A 56-nt DNA strand (CD63T) containing 32-nt DNA aptamer for CD63^[27]^ and 24-nt linker was used to construct DNA nanodevice (Table S1). Three different blockage strategies toward CD63 aptamer in CD63T were rationally designed. The sequence of CD63 aptamer can be blocked in a photocleavable (PC)-group^[28]^ modified hairpin structure, locked by a partially complementary ssDNA containing a PC group (anti19), and locked by two partially complementary ssDNAs containing one PC group (B1 and B2), which were named as HCD, OCD and TCD respectively. Upon UV light irradiation, the PC groups will break apart to open up the hairpin structure of HCD, generate short blocked DNA fragments of OCD and TCD, respectively. As a result, the binding affinities of the residual blocking sequences on the CD63 aptamer were greatly reduced, which facilitate the dissociation of the residual blocking sequences from CD63 aptamer strands in the presence of CD63. The function of CD63 aptamer could be recovered by switching to its active binding motifs.

We first investigated the working efficiency of HCD, OCD and TCD by fluorescence microscopy images of human cervix carcinoma (HeLa) cells. All three responsive lysosome targeting moieties were labeled with 5-carboxyfluorescein (FAM) and lysosomes were stained with Lysotracker red. As shown in Figure S1, confocal fluorescence imaging revealed poor colocalization between three responsive lysosome targeting moieties and lysosomes respectively. The corresponding values of Pearson’s correlation were all around 0.2. After 10-min UV irradiation, the maximum signal overlaps were found between TCD and the lysosomes with Pearson’s correlation coefficient value of 0.7, demonstrating the highest working efficiency to optically controlled lysosomal internalization. Thus, TCD was used for the following experiments.

As a proof-of-concept, the programmed cell death ligand 1 (PD-L1) was chosen as the POI for degradation. The PALTD system was fabricated by using PD-L1T that contains a 35-nt DNA aptamer targeting PD-L1^[29]^ and 24-nt linkers to assemble with TCD (Figure 1a, Table S1). Native polyacrylamide gel electrophoresis PAGE (10 wt%) showed the successful assembly of PALTD systems and gradual dissociation of B2 on the PALTD upon the increasing treating time of UV irradiation (Figure 1b). The PALTD displayed good stability under physiological conditions, since 51% of the system still remained intact after incubation in 10 % fetal bovine serum (FBS) for 24 h (Figure S2). We further investigated the feasibility of PALTD system in the test tube. The B1 were labelled with black hole quencher1 (Q-B1). Due to the fluorescence resonance energy transfer, the fluorescence of FAM on CD63T could be efficiently quenched by the nearby BHQ1 on the Q-PALTD system. The fluorescence intensity of Q-PALTD increased upon light irradiation and incubation with cell lysis solution containing CD63 protein, proving that the PALTD was neither active nor activatable for CD63 binding without light irradiation (Figure 1c). The operation process of Q-PALTD was verified by confocal fluorescence imaging and quantitative analysis of the corresponding signals (Figure 1d).

**Figure 1.**
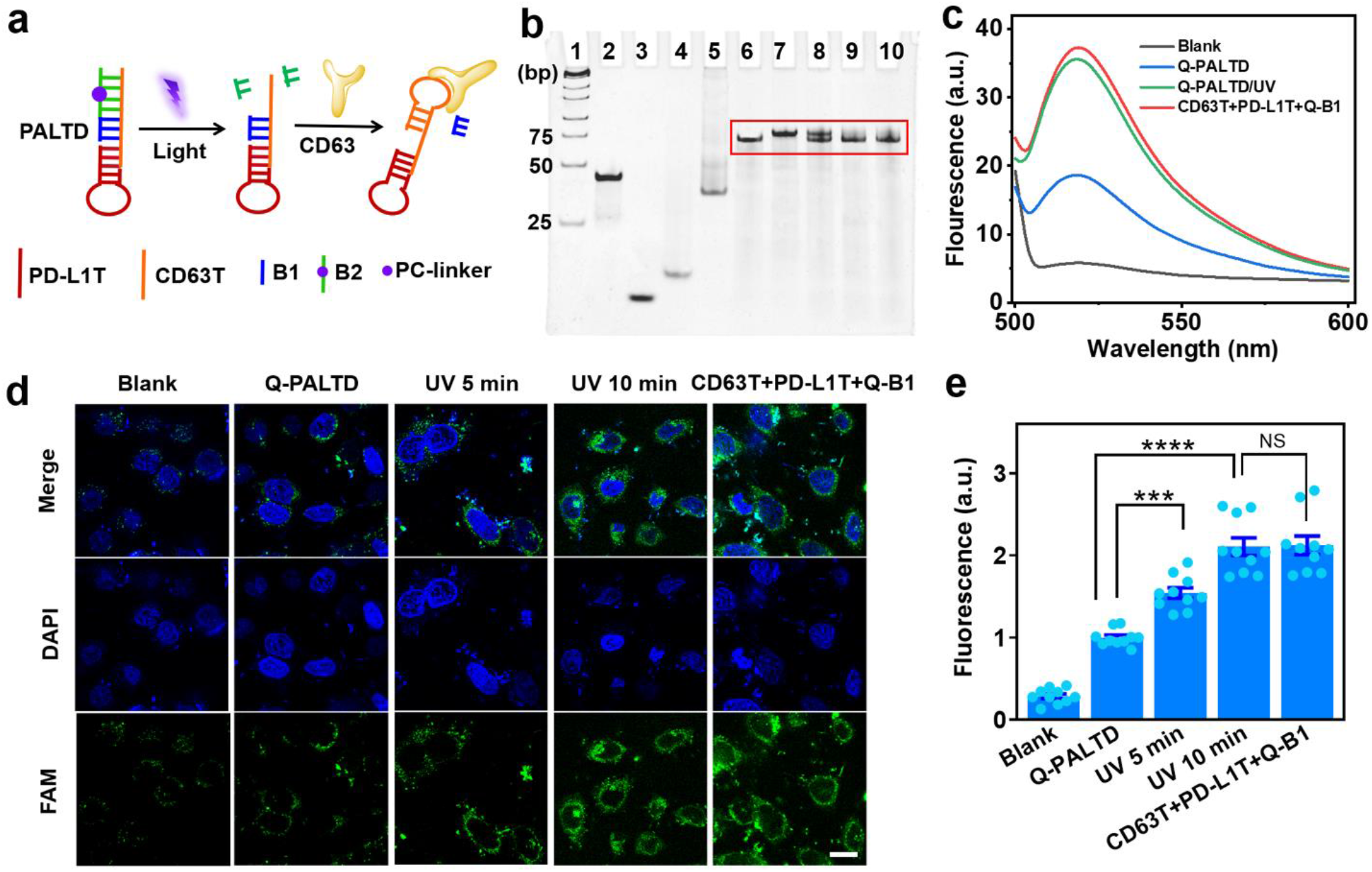
(a) Schematic illustration of the structure and light triggered CD63 binding of PALTD. (b) 10% native polyacrylamide gel analysis of the assembly and UV-triggered dissociation of PALTD. Lanes 1-10 represent DNA ladder marker, CD63T, B1, B2, PD-L1T, CD63T + B1 + PD-L1T, PALTD (CD63T + B1 + B2 + PD-L1T), and PALTD with 5, 10, and 15 min of UV treatments respectively. (c) Fluorescence spectra of different samples incubated with CD63 protein under the excitation wavelength of 480 nm. Black, PBS solution; blue, Q-PALTD (CD63T + Q-B1 + B2 + PD-L1T); green, Q-PALTD treated with 10-min UV irradiation; red, CD63T + Q-B1 + PD-L1T. (d) Confocal fluorescence images of HeLa cells with different treatment, and the fluorescence intensities extracted from corresponding images (e). Scale bar, 20 μm. Shown are mean ± SEM from ten individual cells. ***P < 0.001 (two-tailed Student’s t-test).

We next evaluated the PALTD system for PD-L1 controlled delivery to lysosomes. As shown in Figure S3, upon 10-min irradiation for PALTD before (Light DNA) and after transfection (light cells) to cancer cells, the Cy5 signals of PD-L1 aptamer overlapped well with LysoTracker, with an increasing value of Pearson’s correlation from 0.08 to 0.7 and 0.62 respectively. In contrast, a poor lysosomal accumulation was observed for only treatment of PD-L1 aptamer, verifying the lysosomal-specific internalization. These results demonstrated that PALTD could be selectively activated by the UV light irradiation for PD-L1 delivery. Moreover, confocal imaging suggested that the lysosomal accumulation of PALTD was in an irradiation-time dependent manner and reached saturation at 10 min (Figure S4). These results were in agreement with ELISA analysis of PD-L1 amounts from PALTD-treated cell lysis (Figure S5), which 10 min was selected for the intracellular photoactivation. No significant cytotoxicity was observed in the PALTD system within 20-min irradiation and 24-h incubation (Figure S6). We also wanted to study the kinetics of lysosomal internalization process for PALTD system. After 10-min UV light irradiation, we acquired Cy5 signals of PD-L1 within 2-h incubation in living cells. Increasing the incubation time of PALTD could enhance the uptake of PD-L1 in lysosomes, which the uptake efficiency reached a plateau after 1 h of incubation (Figure S7).

Under the optimal light irradiation and incubation time, the confocal images showed that not only the fluorescence signals of CD63T and PD-L1 overlapped well with each other, but also colocalized well with signals of lysotracker respectively (Figure 2a,b). Further western blot analysis confirmed that the level of PD-L1 proteins sharply decreased after PALTD treatments with 10-min irradiation (Figure 2c), which is consistent with ELISA results Figure S5). The immunofluorescence staining provided the *in-situ* evidences on the elimination of PD-L1 by PALTD in cells (Figure 2d and S8), validating the photoactivated protein degradation.

**Figure 2.**
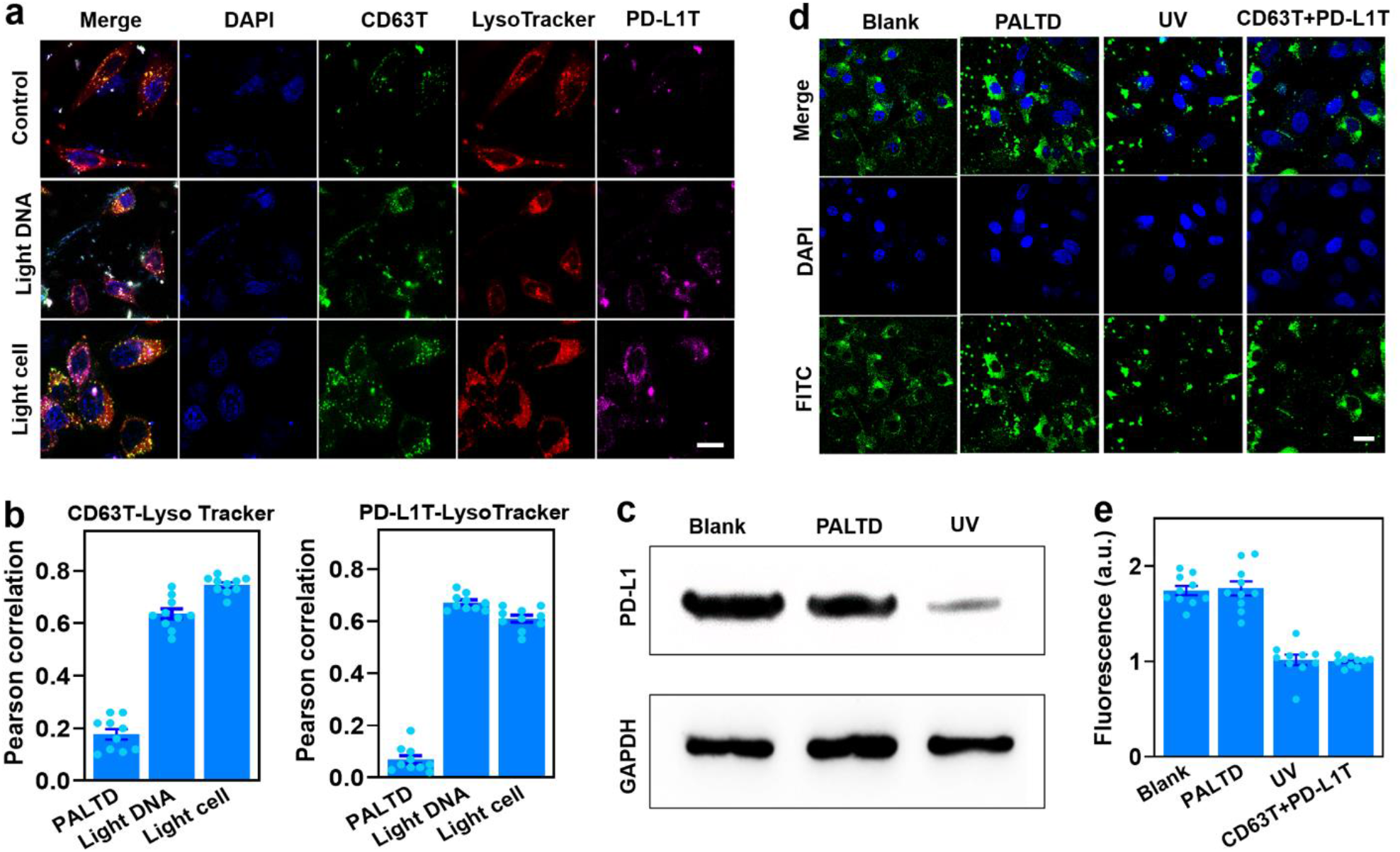
(a) Confocal fluorescence results of PALTD systems with 10 min of UV-treatments before (Light DNA) and after (Light cells) incubation with HeLa cells. Scale bar, 20 μm. (b) Pearson correlation analysis of images (a) by investigating the fluorescence signals of CD63T and PD-L1T with that of Lysotrackers respectively. Shown are mean ± SEM from ten individual cells. (c) PD-L1 amounts after no treatment (blank) and treatment of PALTD with (UV) and without (PALTD) 10 min of UV irradiation. (d) Confocal immunofluorescence staining images of HeLa cells treated with PALTD in the presence (UV) and absence (PALTD) of 10-min UV irradiation, then labeled with PD-L1 antibody (green) and DAPI (blue). Scale bar, 20 mm. (e) The fluorescence intensities extracted from corresponding images (d). Shown are mean ± SEM from ten individual cells.

The DNA-based “AND” logic gate was further designed by using both UV light and glutathione (GSH) as external stimuli to activate lysosomal targeting. The sequence of CD63 aptamer was locked by a fully complementary ssDNA (B3) with rational modification of a PC group and a disulfide bond (Table S1). Either UV or GSH would induce the one-site cleavage of B3, which generates a long DNA fragment still locked the CD63 aptamer. Only both external stimulus could induce two-site cleavage of B3, led to fully dissociation of blocker sequence from CD63 aptamer and the following protein degradation (Figure 3a). Native PAGE (10 wt%) demonstrated the successful assembly and logic operation of CD63/B3 complex (Figure S9).

**Figure 3.**
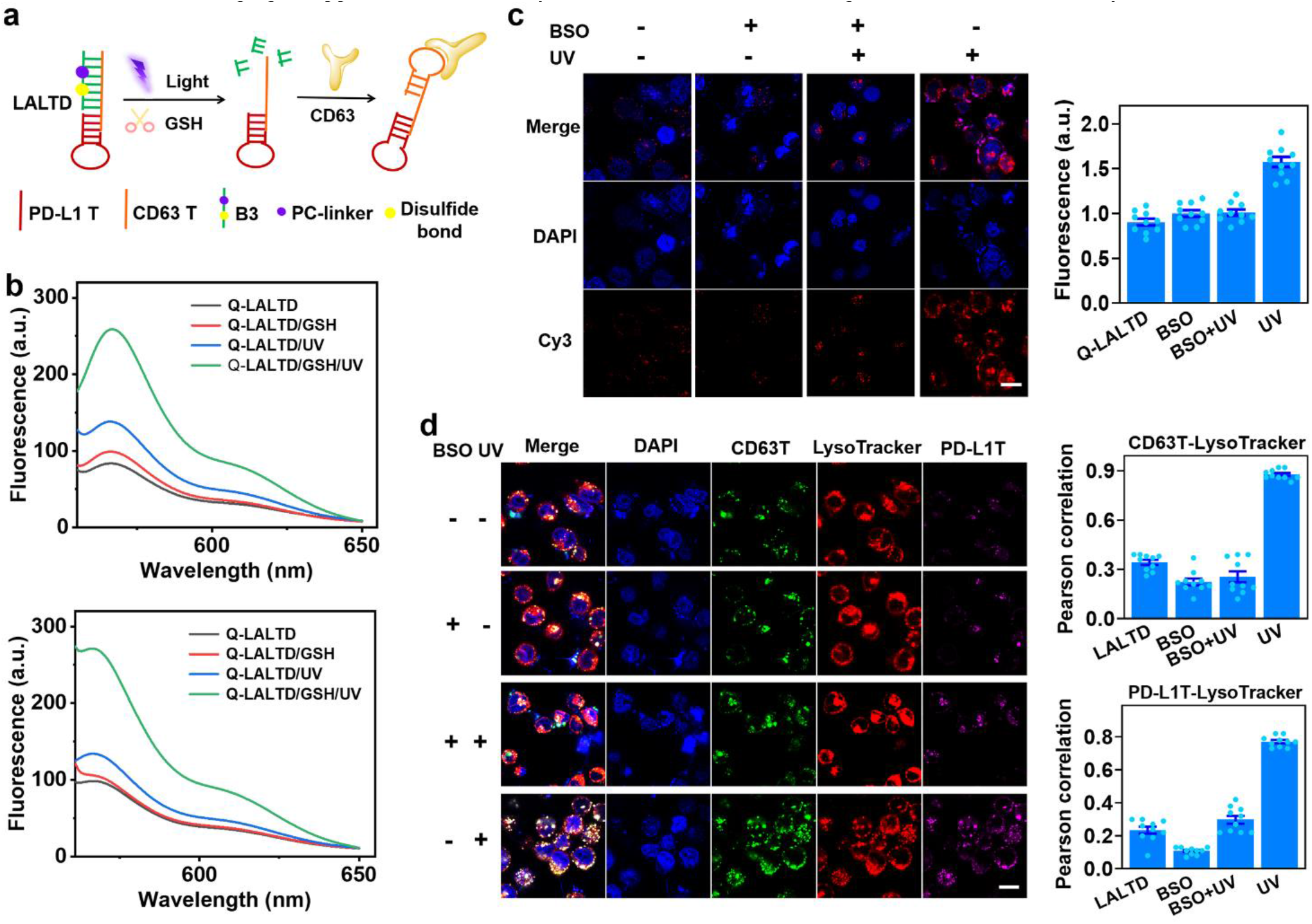
(a) Schematic illustration of the structure and “AND” logic operation of LALTD. (b) Fluorescence spectra of Q-LALTD with different treatments before incubation with CD63 in PBS solution (upper) and lysis solution of HeLa cells (bottom) under the excitation wavelength of 550 nm. (c) Confocal fluorescence images of HeLa cells treated with Q-LALTD under the logic operation by using 100 μM BSO and UV light, as well as fluorescence intensities extracted from the corresponding images. Shown are mean ± SEM from ten individual cells. Scale bar, 20 μm. (d) Colocalization studies on the signal overlapping of lysosomes (lysotracker) with CD63T and PD-L1T respectively. As well as Pearson correlation analysis of the corresponding images by investigating the fluorescence signals of CD63T and PD-L1T with that of Lysotrackers respectively. Shown are mean ± SEM from ten individual cells. Scale bar, 20 μm.

To construct LALTD system, the CD63/B3 complex was connected with PD-L1 aptamer by a long dsDNA of 24 bp as linkers (Table S1). The LALTD demonstrates good stability under physiological conditions (Figure S10). After constructing quenched LALTD (Q-LALTD), the robustness of the DNA-based “AND” logic gate was confirmed by fluorescence analysis in both PBS and extracted protein solutions (Figure 3b). Then, the “AND” logic gate was investigated in living cells by confocal imaging. The overexpressed GSH in cancer cells could be used as the input, which the GSH level was modulated by using buthionine sulfoximine (BSO),^[30]^ an intracellular GSH inhibitor. Only upon UV irradiation without BSO treatment, the increased fluorescence of the Q-LALTD could be observed (Figure 3c).

Then we evaluated the LALTD system for PD-L1 controlled delivery to lysosomes. As shown in Figure 3d, strongest fluorescence signals overlap of both PD-L1 and CD63 aptamers could be observed only when the UV irradiation was applied and without BSO treatments, verifying the logic-gate activated lysosome targeting and internalization of the LALTD system. Further ELISA (Figure S11) and western blot analysis (Figure 4a) confirmed the LALTD system for controlled PD-L1 degradation. The immunofluorescence staining also provided the in-situ evidences on the elimination of PD-L1 by LALTD in HeLa cells (Figure 4b and S12). Moreover, western blot analysis demonstrates that the LALTD also could be applied for PD-L1 protein degradation in 4T1 cells (Figure S13). In order to investigate the universality of the LALTD system, NF-κB subunit p65 (P65) was used as the alternative POI. A 31-nt DNA aptamer^[31]^ targeting P65 was incorporated with CD63T/B3 complex. The controlled P65 degradation was also verified by both immunofluorescence staining (Figure S14a) and western blot analysis (Figure S14b), showing that LALTD system could be used as a general platform for controlled protein degradation.

**Figure 4.**
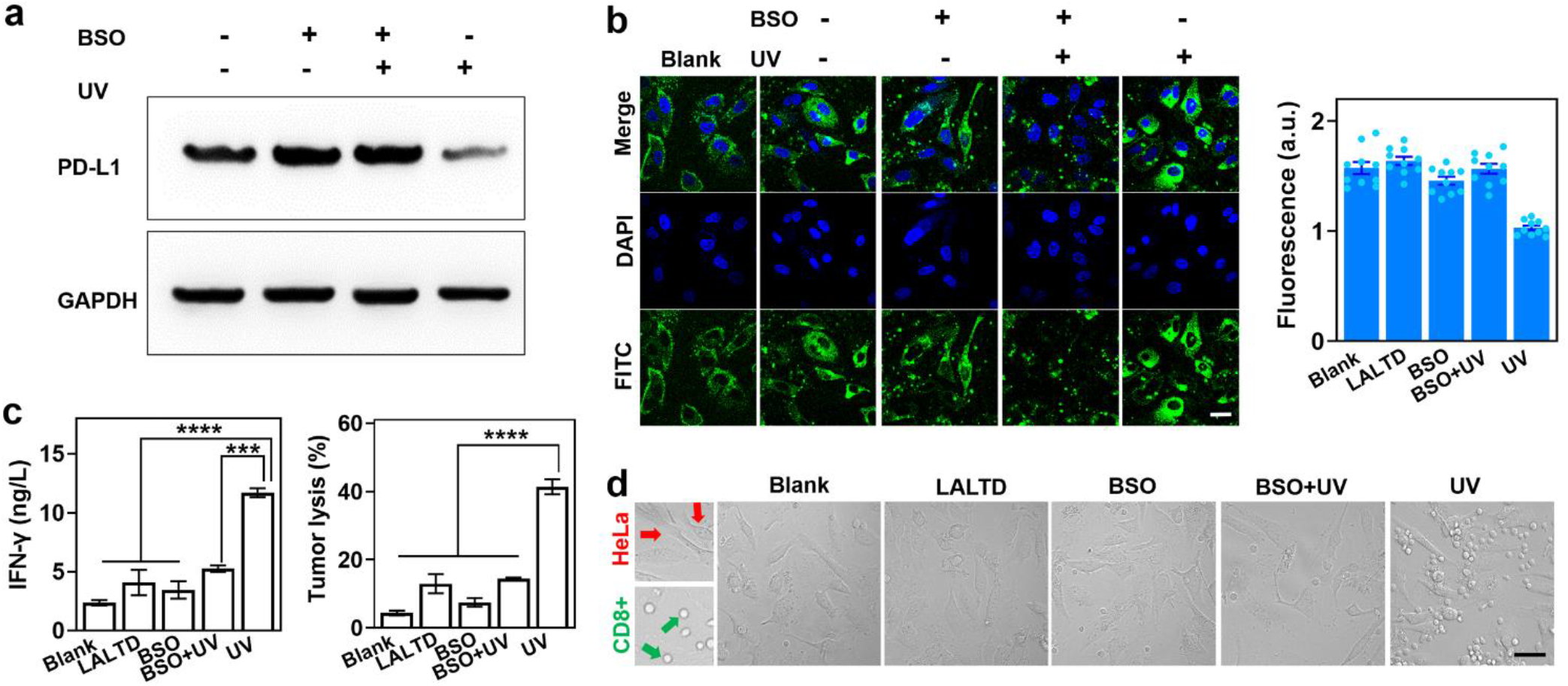
(a) PD-L1 amounts of HeLa cells after LALTD treatments under logic operations by 100 μM BSO and UV light. (b) Confocal immunofluorescence staining images of Hela cells treated with LALTD under logic operations by 100 μM BSO and UV light, then labeled with PD-L1 antibody (green) and DAPI (blue), and fluorescence intensities extracted from the corresponding images. Shown are mean ± SEM from ten individual cells. Scale bar, 20 μm. (c) Quantification of IFN-γ and tumor lysis percentage by CD8+ T-cells by using LALTD under logic operations. Shown are mean ± SEM (n = 3). (d) Confocal microscopic images of HeLa cells treated with LALTD under logic operation, and then co-incubated with CD8+ T-cells for 24 h. Scale bar, 25 μm.

T cells with PD-1 ligand could recognize cancer cells with PD-L1 receptor, followed by efficient inhibition of interferon-γ (IFN-γ) secretion and tumor cell lysis.[32] Thus, we studied deeper on the relation between PD-L1 degradation and CD8+ T cell-mediated antitumor immunity by LALTD system. The T cells were incubated with PALTD treated HeLa cells under different logic operations. The highest levels of IFN-γ and tumor lysis percentage could be achieved only by UV irradiation without BSO treatments (Figure 4c). Confocal images verified the strongest T cells recognition toward cancer cells by the (1, 1) input of LALTD system (Figure 4d), suggested that effective blockage of PD-1/PD-L1 interactions by precisely controlled PD-L1 degradation.

Besides, the LALTD system can be easily labeled with cell targeting moieties as well, such as folate (Table S1), for strengthening the cell targeting and uptake. Confocal fluorescence imaging revealed the successful internalization and logic operation of folate-LALTD in living cells (Figure S15), which the controlled PD-L1 degradation was confirmed by the immunofluorescence staining (Figure S16a) and western blot analysis (Figure S16b)

In summary, we have constructed a general platform for controlled proteins degradation via activatable lysosome targeting DNA nanodevices in living cells. The locked CD63 aptamer on the PALTD and LALTD systems could be activated by UV irradiation and UV/GSH based “AND” logic gate operation for efficient targeting PD-L1 into lysosomes. The ELISA, western blot analysis and immunofluorescence staining confirmed the successfully efficient degradation of target protein. The LALTD systems could further manipulate T cell-mediated antitumor immunity by logic operation. By changing the aptamer of POI, the modular designed system demonstrated robust universality for precisely controlled protein degradation.

## Acknowledgements

The authors gratefully acknowledge the National Natural Science Foundation of China (22104058, 22174066), and the Program of Jiangsu Specially-Appointed Professor.

