## SUPPORTING INFORMATION for "Logic Gate Activated Lysosome Targeting DNA Nanodevice for Controlled Proteins Degradation"

Supporting Information
©Wiley-VCH 2021
69451 Weinheim, Germany

### Logic Gate Activated Lysosome Targeting DNA Nanodevice for Controlled Proteins Degradation

Yuzhe Shang, Longyi Zhu, Yang Xiao, Songyuan Du, Ruoyang Ji, Bin Li, Jialiang Chen, Shengyuan Deng,* Kewei Ren*

**Abstract:** Targeted protein degradation (TPD) is a powerful technique for regulation of protein homeostasis. Current TPD mainly focus on the therapeutical consequences rather than the operation processes of the molecular tools. Herein, we construct a platform for precisely manipulate the protein degradation by activatable lysosome targeting DNA nanodevices. In the design, a lysosome-targeting CD63 aptamer is locked by the single-stranded DNA with a photocleavable group and a disulfide bond. This locked CD63 aptamer is connected with the aptamer targeting the protein of interest via double-stranded DNA linkages to form the logic-gate activated lysosome targeting DNA nanodevice (LALTD). With the UV light and endogenous GSH as inputs, AND logic-gate is built to efficiently manipulate the protein delivery processes by LALTD. The protein of interest could be degraded via efficient lysosome hydrolysis. Further studies showed that the logic-gate operation could be used for modulating the T cell-mediated antitumor immunity. The designed activatable lysosome targeting DNA nanodevices exhibits good stability, controllability, and programmability, providing a new prospect for accurate protein degradation.

**DOI: 10.1002/anie.2021XXXXX**

#### Table of Contents

**Experimental Procedures**......S2
**Supporting Table S1**......S3
**Supporting Figures S1-S16**......S4

#### Experimental Procedures

**Reagents.** All the chemicals were purchased from Sigma unless otherwise noted. Commercial reagents were used without further purification. LysoTracker red, 4',6-Diamidino-2-Phenylindole (DAPI), yeast tRNA, bovine serum albumin (BSA) and SYBR gold were purchased from Invitrogen (Carlsbad, CA, USA). Fetal bovine serum (FBS), glucose, agarose, 40% polyacrylamide, TEMED, APS and all of the DNA were from by Sangon Biotech Co., Ltd (Shanghai, China). Dulbecco's PBS (DPBS) was obtained from GlpBio Technology Inc (Montclair, CA, USA). The detailed sequences were listed in Table S1. Washing buffer was prepared by mixing 4.5 g/L glucose and 5 mM $MgCl_2$ in Dulbecco's PBS with calcium chloride and magnesium chloride. Binding buffer was prepared by adding yeast tRNA (0.1 mg/mL) and BSA (1 mg/mL) in washing buffer to reduce non-specific binding. All aqueous solutions were prepared using ultrapure water (18.2 MΩ·cm, Milli-Q, Millipore). PD-L1 antibody (13684T), GAPDH antibody (5174T), NF-κB p65 antibody(8242T) and Anti-rabbit IgG, HRP-linked Antibody (7074P2) were from Cell signaling. Goat Anti-Rabbit IgG (FITC) was from YIFEIXUE BIOTECH CO., LTD. (Nanjing, China).

**Apparatus.** The concentrations of nucleic acids were all measured using a NanoDrop One UV-Vis spectrophotometer. The gel electrophoresis was performed on a Tanon EPS-300 Electrophoresis Analyser (Tanon Science & Technology Company, China) and imaged on Bio-rad ChemDoc XRS (Bio-Rad, USA). All the *in vitro* fluorescence measurements were measured on an F-7000 spectrometer (HITACHI, Japan). All the intracellular images were taken by a Nikon A1& SIM-S&STORM super resolution microscope (Tokyo, Japan). MTT and ELISA assays were measured with a Safire microplate Analyzer (Molecular Devices, America).

**Cell culture.** Human cervix carcinoma (HeLa) cells lines (Procell Life Science & Technology, Wuhan, China) were cultured in Dulbecco's modified Eagle's medium (DMEM) supplemented with 10% fetal bovine serum (FBS), 100 mg/mL streptomycin and 100 U/mL penicillin-streptomycin at 37 ºC in a humidified incubator containing 5% $CO_2$ and 95% air. Short tandem repeats (STR) profiling and mycoplasma testing were conducted for each cell line before use. Cell numbers were determined with a Petroff-Hausser cell counter (USA).

**Activatable DNA nanodevices construction.** ssDNA strands CD63T, B1 (or Q-B1), B2 and PD-L1T were mixed together to 4 μM in 50 μL DPBS containg 20 mM $MgCl_2$. After heating at 95 ºC for 5 min, the mixture was slowly cooled to 25 ºC at a rate of 1 ºC / min to obtain the photo-activatable DNA nanodevices (PALTD). The logic-gate activated lysosome targeting DNA nanodevice (LALTD or Q-LALTD) was obtained by using CD63T (or CD63L or Folate-CD63T), PD-L1-T (or P65T) and B3 (or Q-B3) following the same approach.

**Polyacrylamide gel electrophoresis analysis (PAGE).** Native polyacrylamide gel (10%) was prepared using 1 × TBE buffer. The loading samples were prepared by mixing 7.5 μL DNA samples and 1.5 μL 6 × loading buffer, and placed still for 3 min before injected into polyacrylamide gel. The gel electrophoresis was run at 100 V for 50 min in 1 × TBE buffer, stained with 1 × SYBR Gold, and scanned with a Molecular Imager Gel Doc XR.

**Serum stability of activatable DNA nanodevices.** 1 μM of PALTD or LALTD were incubated with 10% FBS for different time (1-24 h). 10% native polyacrylamide gel in 1 × TBE buffer was run at 120 V for 90 min and stained with 1 × SYBR Gold staining solution.

**3-(4,5-dimethylthiazol-2-yl)-2,5-diphenyltetrazolium bromide (MTT) assay.** Hela cells were cultured within 96-well plates at a density of $1\times10^4$ cells/chamber for 24 h. After discarding the medium, the cells were washed with washing buffer and incubated with or without the DNA nanodevices transfected with Lipofectamine® 2000 transfection reagent. After 24 hour-incubation, the cells were washed twice with washing buffer and treated with or without 365-nm light (3 mW/cm$^2$) for different times. These treated cells were continuously cultured for 24 h and washed with washing buffer for two times. Then 50 μL of 5 mg/ml MTT solution was added and incubated for 4 h. After removing the remaining MTT solution, 100 μL of dimethylsulphoxide was added and the plate was shaken for 10 min, then the optical density at a wavelength of 490 nm was measured with a Safire microplate Analyzer.

**Confocal fluorescence imaging data analysis.** $1\times10^4$ HeLa cells were seeded in a confocal dish for 24 h at 37 ºC, then transfected with or without 1 μM DNA nanodevice using a Lipofectamine® 2000 transfection reagent (Thermo Fisher Scientific) for 24 h. Then the cells were washed twice with washing buffer and treated with or without 365-nm light (3 mW/cm$^2$) for 10 min. After 12 hour-incubation, the cells were washed twice with washing buffer and stained with 5 mg/mL of DAPI for 15 min for imaging.

For the lysosomal colocalization experiment, $1\times10^4$ HeLa cells were seeded in a confocal dish for 24 h at 37 ºC, then transfected with or without DNA nanodevices using a Lipofectamine® 2000 transfection reagent (Thermo Fisher Scientific) (or incubated with folate-LALTD) for 24 h. Then the cells were washed twice with washing buffer and treated with or without 365-nm light (3 mW/cm$^2$) for 10 min. After 12 hour-incubation, the cells were washed twice with washing buffer, as well as stained with 50 nM of LysoTracker Red and 5 mg/mL of DAPI for 15 min for imaging.

All the fluorescence images were collected with Nikon A1& SIM-S&STORM super resolution microscope. LysoTracker red, Cy5, FAM and DAPI were excited with 561 nm, 640 nm, 488 nm and 405 nm lasers, respectively. A 100× oil immersion objective was used for imaging cells. Image analysis was performed with a NiS-Elements AR Analysis software. Data analysis and fitting was done using the Origin and GraphPad Prism software.

**ELISA for protein detection.** HeLa cells were seeded into 24-well plate at 5 × $10^5$ per well and incubated for 24 h at 37 ºC. After washing with DPBS, the cells were incubated with or without serial concentrations of the DNA probes transfected with Lipofectamine® 2000 transfection reagent. After 24 hour-incubation, the cells were washed twice with washing buffer and treated with or without 365-nm light (3 mW/$cm^2$) for 10 min. These treated cells were continuously cultured for 24 h and washed with DPBS for two times, then lysed with radioimmunoprecipitation assay (RIPA) buffer on ice for 30 min. After the cells were centrifuged at 12,000 g for 10 min, the supernatant was collected and the PD-L1 protein concentration was measured by ELISA.

**Western blot**. To detect the effect PALTD, LALTD or folate-LALTD on the level of PD-L1 protein (or transcription factor p65 protein, P65), HeLa cells (or 4T1 cells) were treated without (Blank) or with DNA nanodevice transfected with Lipofectamine® 2000 transfection reagent for 24 h. Then the cells were washed twice with washing buffer and treated with or without 365-nm light (3 mW/$cm^2$) for 10 min. After 24-h incubation, cells were harvested and followed by adding SDS loading buffer for western blot. The levels of PD-L1, P65 protein and GAPDH were analyzed by immunoblotting using antibodies against PD-L1, P65 and GAPDH, respectively.

**Immunofluorescence staining**. HeLa cells treated without (Blank) or with PALTD or LALTD transfected with Lipofectamine® 2000 transfection reagent, or folate-LALTD respectively for 24 h. Then the cells were treated with or without 365-nm light (3 mW/cm2) for 10 min. After 24-h incubation, the cells were fixed by 4% paraformaldehyde for 10 minutes. After preblocking for 1 h in PBS with 10% FBS (v/v) and 5% BSA bovine serum albumin (w/v), cells were incubated with PD-L1 (or P65) antibody for 2 h at room temperature and then with FITC goat Anti-Rabbit IgG for 1 h at room temperature, followed by incubation with 5 mg/mL of DAPI for 15 min for imaging. The cells were observed under Nikon A1& SIM-S&STORM super resolution microscope. FITC and DAPI were excited with 488 nm and 405 nm lasers, respectively.

**T-cell cytotoxicity assay.** HeLa cells treated without (Blank) or with LALTD transfected with Lipofectamine® 2000 transfection reagent for 24 h. Then the cells were treated with or without 365-nm light (3 mW/cm2) for 10 min. After 24-h incubation, the HeLa cells were co-cultured with activated CD8+ T-cell (Milestone® Biotechnologies, Shanghai, China) for 24 h. Then HeLa cells co-incubated with CD8+ T-cells were observed under microscope, and IFN-γ in the cell culture medium was measured using an IFN-γ ELISA Kit (YIFEIXUE BIOTECH CO., LTD., Nanjing, China). The viability of HeLa cells after co-culture was assessed using the CCK-8 assay.

**Table S1.** DNA sequences used in the experiments.

| Names | Sequences from 5’ to 3’ |
|---|---|
| CD63 Aptamer (CD63) | CACCCCACCTCGCTCCCGTGACACTAATGCTA-FAM |
| HCD | GTGTCACGGIPClinkerCACCCCACCTCGCTCCCGTGACACTAATGCTATTTTTTTTTTTTTTTTTTTTTTTT-FAM |
| anti19 | AGTGTCACGIPClinkerGGAGCGAGGT |
| CD63T[a] | CACCCCACCTCGCTCCCGTGACACTAATGCI6FAMdTAGTTCTATGTTTCTTTGTATCTTTG |
| Blocker 1 (B1) | GCATTAGTGTCA |
| Blocker 2 (B2) | CGGGAGCGAIPClinkerGGTGGGGTG |
| Q-B1 | BHQ1-GCATTAGTGTCA |
| Blocker 3 (B3) | TAGCATTAGTGIPClinkerTCACGGGAGCGSH-SHAGGTGGGGTG |
| CD63L | CACCCCACCTCGCTCCCGI6Cy3dTGACACTAATGCTAGTTCTATGTTTCTTTGTATCTTTG |
| Q-B3 | TAGCATTAGTGIPClinkerIBHQ1dTCACGGGAGCGSH-SHAGGTGGGGTG |
| PD-L1T[b] | Cy5-TACAGGTTCTGGGGGGTGGGTGGGGAACCTGTTTTCAAAGATACAAAGAAACATAGAAC |
| P65T | TGGGGACTTTCCAGTTTCTGGAAAGTCCCCATTCAAAGATACAAAGAAACATAGAAC |
| Folate-CD63T | Folate-CACCCCACCTCGCTCCCGTGACACTAATGCI6FAMdTAGTTCTATGTTTCTTTGTATCTTTG |

[a] The base-pairing region was labelled with purple colour. [b] The aptamer sequence targeting PD-L1 was underlined.

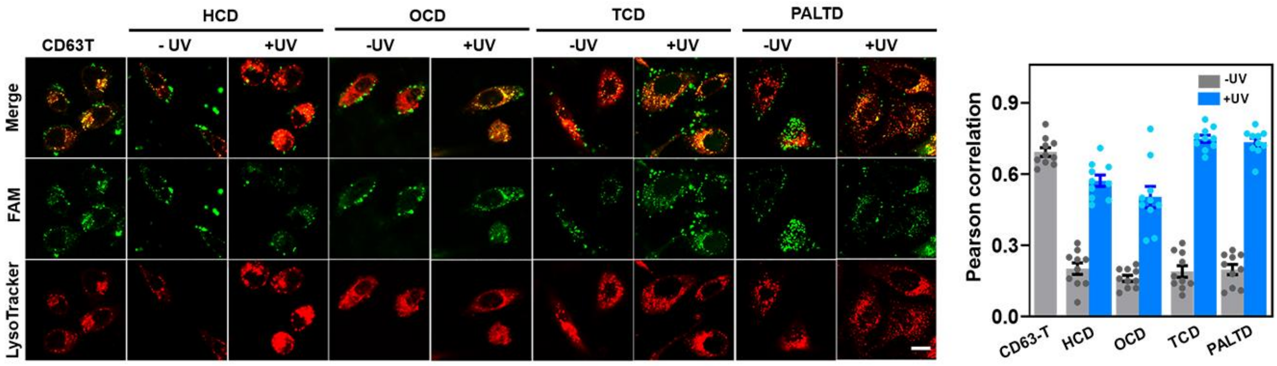


**Figure S1.** Confocal fluorescence images of HeLa cells incubated with HCD, OCD, TCD and PALTD with and without 10 min of UV irradiations. Scale bar, 20 μm. The Pearson correlations were measured by using the FAM signals of CD63T and signals of lysotracker in the corresponding images. Shown are mean ± SEM from ten individual cells.

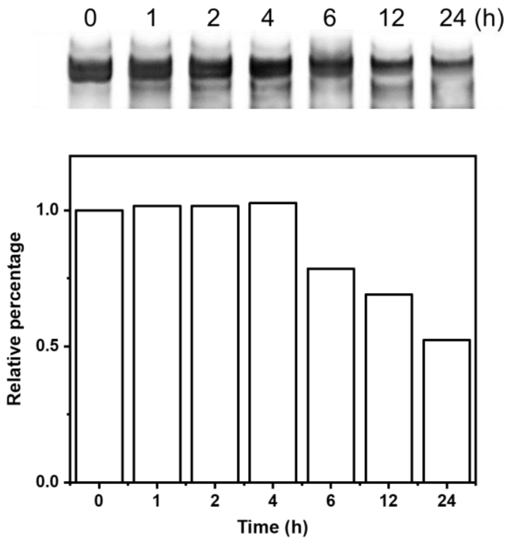


**Figure S2.** The stability study of PALTD systems in 10 % fetal bovine serum (FBS) for 24 h by 10% denature PAGE analysis.

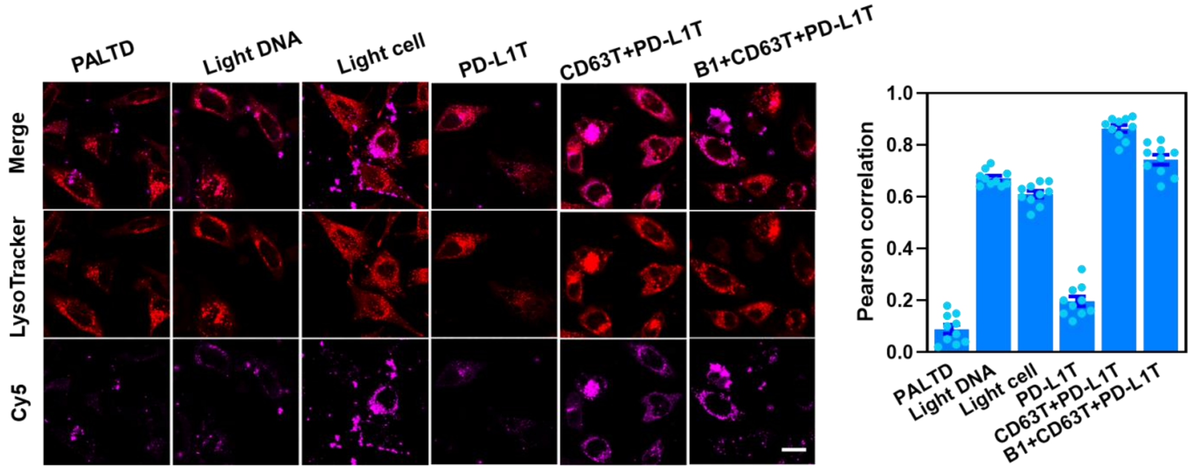


**Figure S3.** Confocal fluorescence results of PALTD systems with 10 min of UV-treatments before (Light DNA) and after (Light cells) incubation with Hela cells, as well as control experiments by using PD-L1T, CD63T + PD-L1T, B1 + CD-T + PD-L1T respectively. The Pearson correlations were measured by using the Cy5 signals of PD-L1T and signals of lysotracker in the corresponding images. Scale bar, 20 μm. Shown are mean ± SEM from ten individual cells.

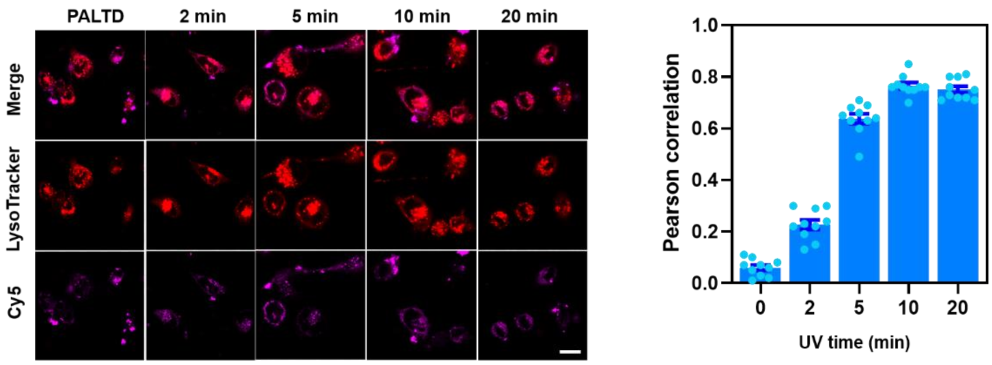


**Figure S4.** Confocal fluorescence results of PALTD treated Hela cells under different time of UV irradiations. The Pearson correlations were measured by using the Cy5 signals of PD-L1T and signals of lysotracker in the corresponding images. Scale bar, 20 μm. Shown are mean ± SEM from ten individual cells.

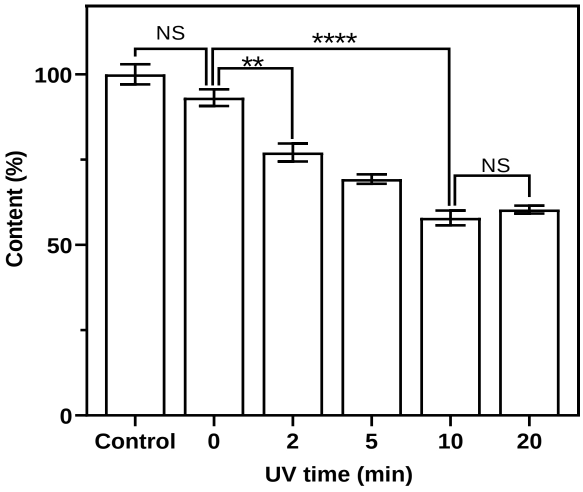


**Figure S5.** ELISA analysis on the PD-L1 Levels of Hela cells (control), and PALTD treated Hela cells with different irradiation time. Shown are mean ± SEM (n = 3).

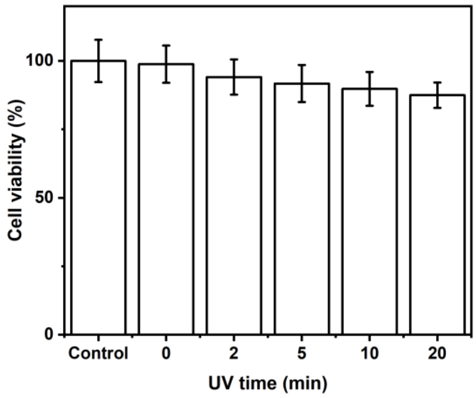


**Figure S6.** Cell viability study of PALTD systems under different time of UV irradiations. Shown are mean ± SEM (n = 3).

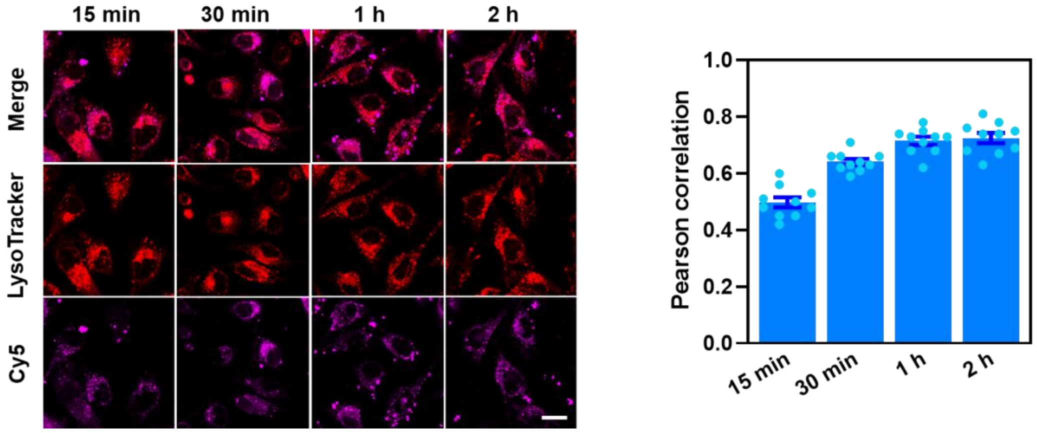


**Figure S7.** Confocal fluorescence results of PALTD treated Hela cells for different incubation time. The Pearson correlations were measured by using the Cy5 signals of PD-L1T and signals of lysotracker in the corresponding images. Shown are mean ± SEM from ten individual cells.

.

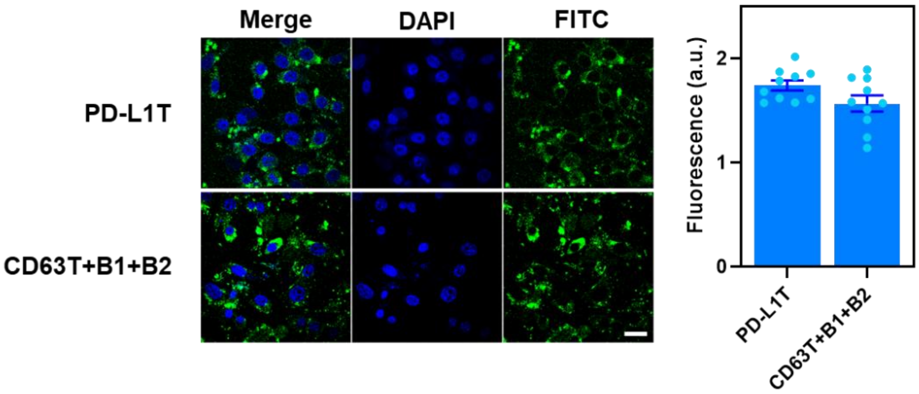


**Figure S8.** Confocal immunofluorescence staining images of Hela cells treated with PD-L1T and CD63T + B1 + B2, then labeled with PD-L1T antibody (green) and DAPI (blue), as well as the fluorescence intensities extracted from corresponding images. Scale bar, 20 μm. Shown are mean ± SEM from ten individual cells.

.

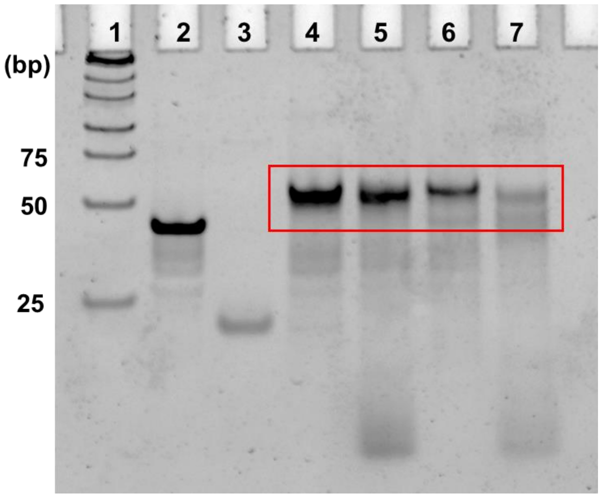


**Figure S9.** 10% denature PAGE analysis of the assembly and logic operation of LALTD. Lanes 1-7 represent DNA ladder marker (25-500 bp), CD63L, B3, CD63L + B3, CD63L + B3 after 10 min of UV irradiation, CD63L + B3 after adding 10 mM of GSH, and CD63L + B3 by both 10-min UV and 10-mM GSH treatments.

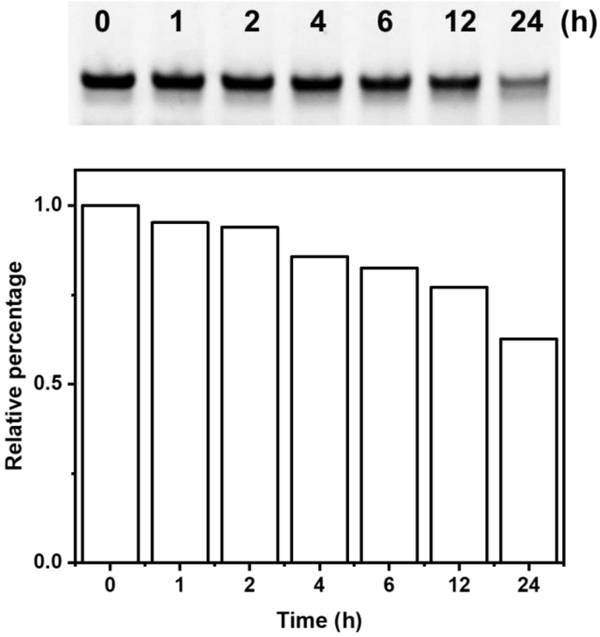


**Figure S10.** The stability study of LALTD systems in 10 % fetal bovine serum (FBS) within 24 h by 10% denature PAGE analysis.

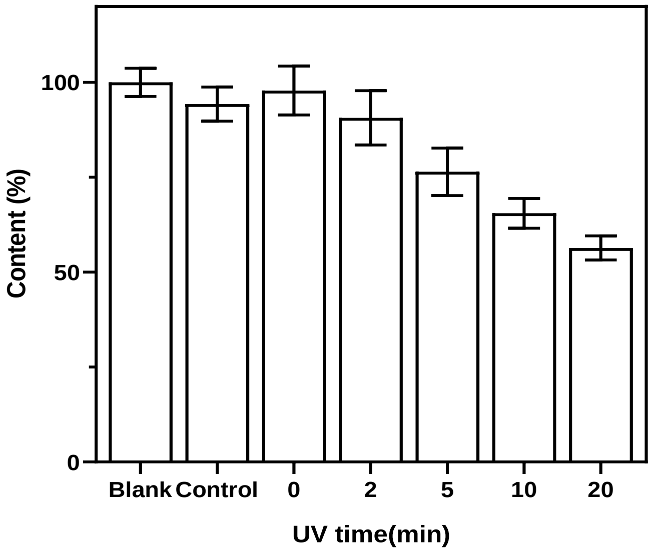


**Figure S11.** ELISA analysis on the PD-L1 Levels of Hela cells (Blank), LALTD treated Hela cells with different UV irradiation time, or with 10-min UV irradiation and 100 μM BSO (Control). Shown are mean ± SEM (n = 3).

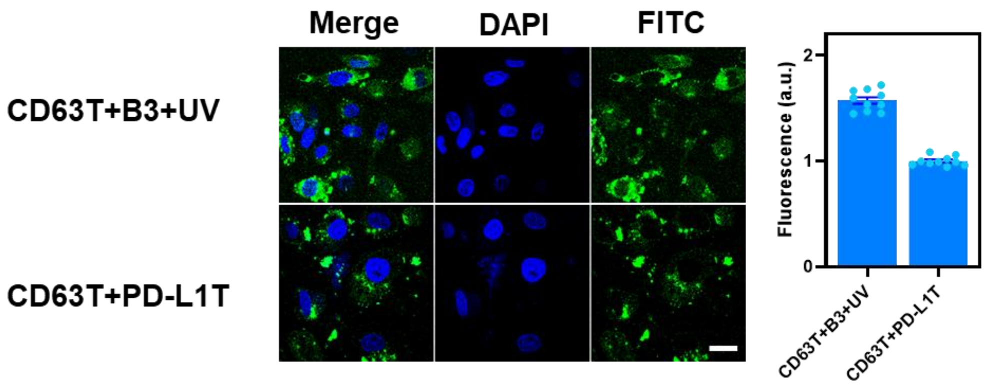


**Figure S12.** Confocal immunofluorescence staining images of Hela cells treated with CD63T + B3 under 10-min UV irradiation and CD63T + PD-L1T, then labeled with PD-L1T antibody (green) and DAPI (blue), as well as the fluorescence intensities extracted from corresponding images. Scale bar, 20 μm. Shown are mean ± SEM from ten individual cells.

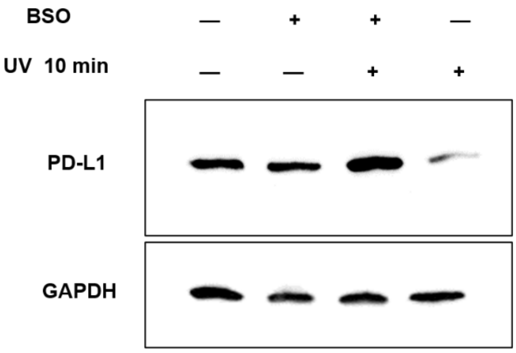


**Figure S13.** PD-L1 amounts of 4T1 cells after LALTD treatment under logic operations by 100 μM BSO and UV light.

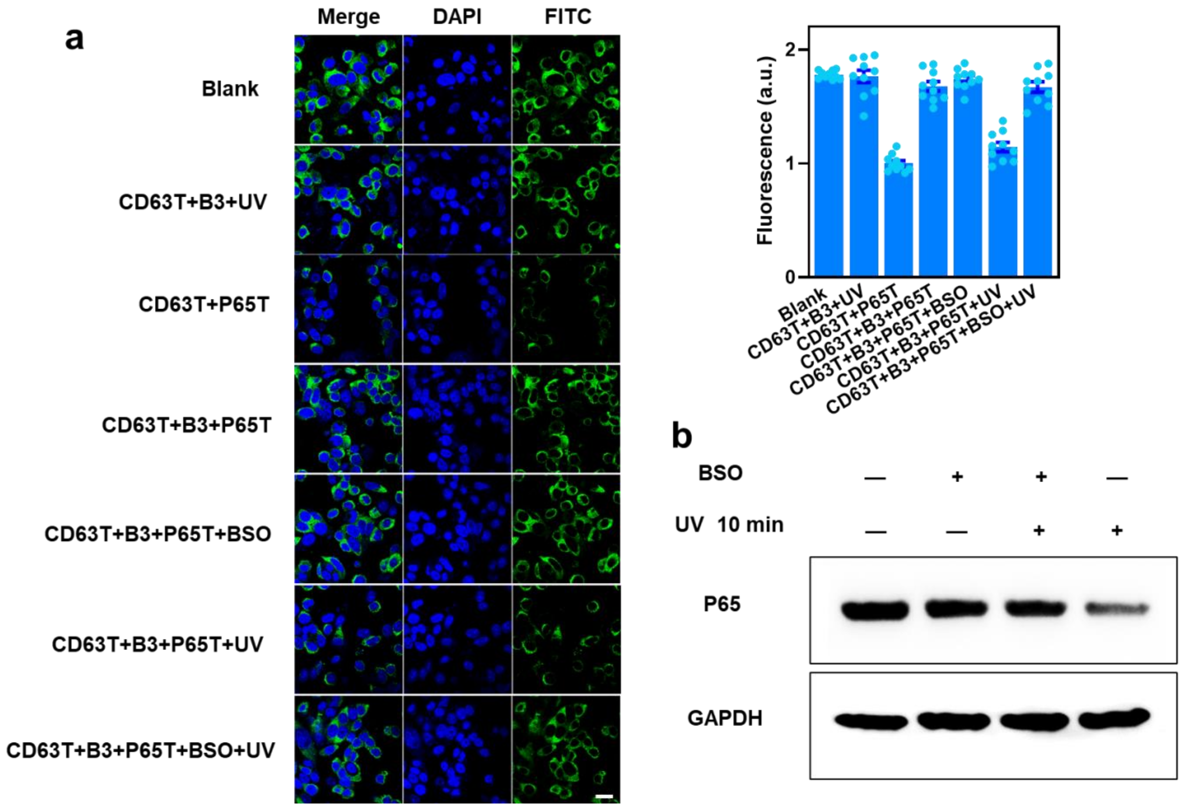


**Figure S14.** (a) Confocal immunofluorescence staining images of Hela cells with different treatments, then labeled with P65T antibody (green) and DAPI (blue), as well as the fluorescence intensities extracted from corresponding images. Scale bar, 20 μm. Shown are mean ± SEM from ten individual cells. (b) P65 amounts after LALTD treatment under logic operations by 100 μM BSO and UV light.

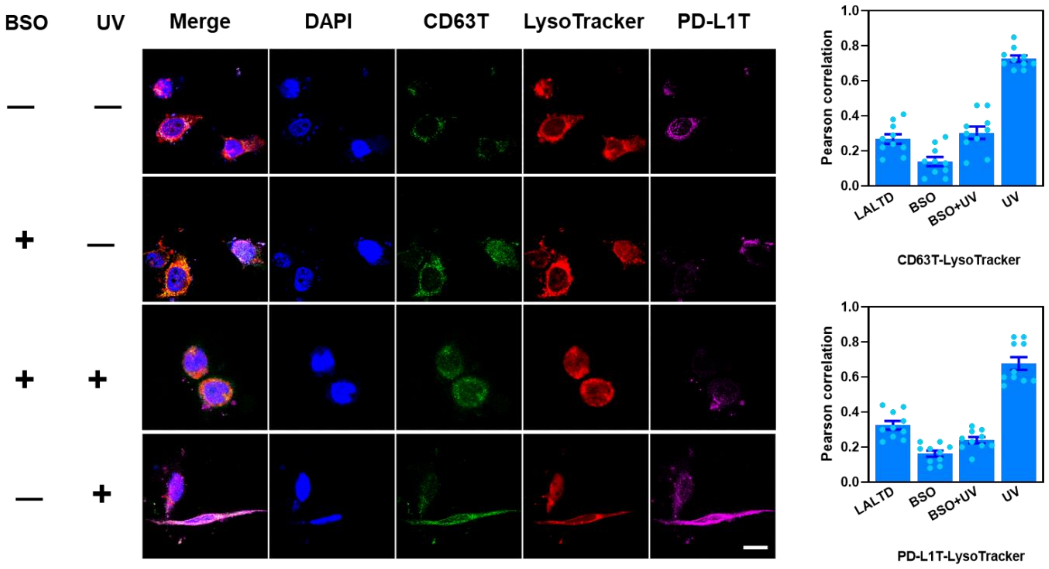


**Figure S15.** Colocalization studies on the signal overlapping of lysosomes (lysotracker) with dual signals on the folate-LALTD systems (Folate-CD63T + B1 + B2 PD-L1T) respectively. Scale bar, 20 μm. Pearson correlation analysis by investigating the fluorescence signals of CD63T and PD-L1T on the folate-LALTD systems with that of Lysotrackers respectively. Shown are mean ± SEM from ten individual cells.

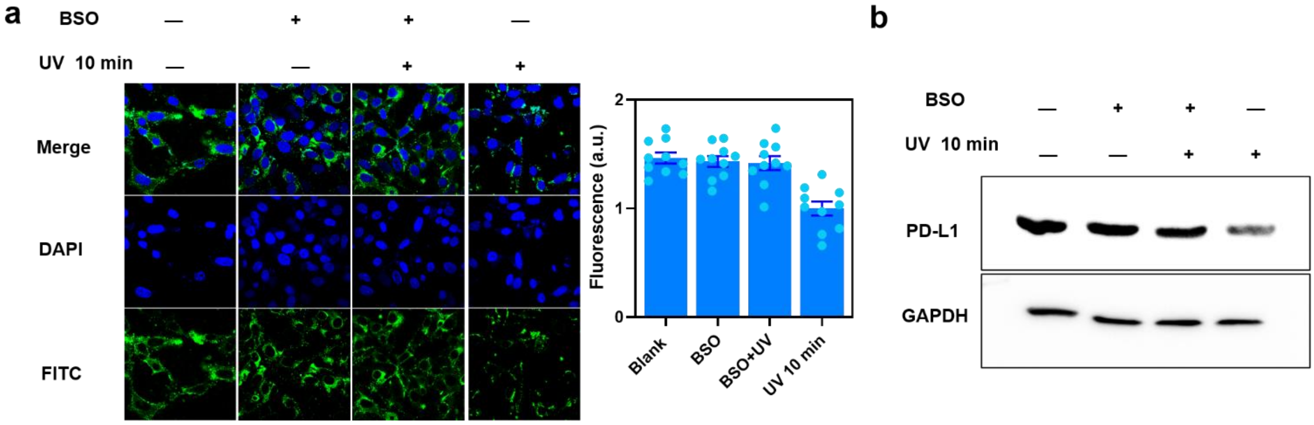


**Figure S16.** (a) Confocal immunofluorescence staining images of Hela cells treated with folate-LALTD under logic operations by 100 μM BSO and UV light, then labelled with PD-L1 antibody (green) and DAPI (blue), and fluorescence intensities extracted from the corresponding images. Shown are mean ± SEM from ten individual cells. Scale bar, 20 μm. (b) PD-L1 amounts of HeLa cells after folate-LALTD treatment under logic operations by 100 μM BSO and UV light.
